# Nitrate regulates anchor root development

**DOI:** 10.64898/2026.08.31.748355

**Authors:** Suresh Damodaran, Michel Ruiz Rosquete, Natalie Gonzalez, Wolfgang Busch, Lucia C. Strader

## Abstract

Nitrogen is a critical nutrient necessary for plant growth and survival. Plasticity in root architecture helps adapt to soil nitrogen levels for optimal nitrogen uptake; the nitrate form of soil nitrogen is a major modulator of root architecture. Although details of nitrate-regulated primary and lateral root growth are known, nitrate-regulated formation of anchor roots, which arise from the collet, is not understood. In this work, we uncover a role for nitrate in the regulation of anchor root formation. We find that cytokinin inhibits anchor root formation with rising nitrate. These cytokinin effects on anchor root formation rely on regulated indole-3-butyric acid (IBA) to indole-3-acetic acid (IAA) conversion. These data point toward a mechanism by which nitrate controls a previously underappreciated aspect of nitrate-dependent root architecture driven by anchor roots.

**Summary Statement:** Elevated nitrate levels promote anchor root formation to alter root growth. In Arabidopsis, cytokinin restricts anchor root formation at elevated nitrate levels through indole-3-butyric acid (IBA) derived auxin.

## 1 Introduction

Root system architecture is typically evaluated based on the primary and lateral roots contributing to the spatial configuration. In particular, the frequency, length and branching angle of lateral roots along primary roots determine the root system architecture and exploration of soil resources. Anchor roots, which emerge from sites at the root-shoot junction, are an underappreciated contributor to dicot root system architecture. Anchor root formation is generally promoted in response to the loss of the primary root meristem, and auxin is essential for their formation (Lucas et al., 2011, Jia et al., 2019, Bai et al., 2020). Further, anchor root formation is promoted by the endogenous carotenoid-derived compound anchorene and low N (Jia et al., 2019). The contribution of anchor roots to the overall architecture of the root system and their role in nutrient acquisition have not been fully explored.

Anchor roots emerge from the collet, which is the junction between the hypocotyl and roots in eudicots such as Arabidopsis. These are distinct from adventitious roots, which can emerge from any non-root organ, including the hypocotyl, stem, leaf, or callus (Bellini et al., 2014). In addition, the founder cells of anchor roots are pericycle cells in the collet, as opposed to adventitious roots which arise from a wider range of founder cells including, the cambium, vascular parenchyma, or dedifferentiated cells, in addition to the pericycle. Unlike the loosely patterned adventitious roots, anchor roots typically emerge oppose one another in the collet (Jia et al., 2019). Finally, these organs are driven by endogenous and metabolite signaling, rather than the largely wound- or stress-induced adventitious roots, suggesting that anchor roots are distinct from adventitious roots.

The collet is specified during embryo development (Scheres et al., 1994). This region is anatomically distinct from the primary root and hypocotyl, as it develops an additional layer of cortex and all epidermal cells develop root hairs. Moreover, collet epidermal hairs may be involved in nutrient sensing at the soil surface (Parsons, 2009). Given the combination of nutrient sensing by collet root hairs and the variable emergence of anchor roots, we hypothesize that this root region plays a key role in integrating nutrient signals and root architecture. Indeed, anchor root formation has previously been shown to respond to nitrate and phosphate levels (Jia et al., 2019).

Developmental plasticity enables plants to adapt their root growth in response to nutrient and water availability. Nitrogen is a limiting nutrient for plant growth and a major determinant of adaptive root growth (Crawford and Forde, 2002). In well-aerated soil, nitrate is the primary nitrogen source of nitrogen; however, due to its solubility and high mobility, uncaptured nitrate leaches deeper into the soil or into nearby water bodies. Thus, there exists spatial variation in soil nitrate availability, requiring plants to adapt their root architecture to maximize nitrate acquisition (Crawford and Glass, 1998). Understanding nitrogen-regulated root development could provide tools for maximizing efficient plant usage of soil nitrogen (Lamig et al., 2022).

Nitrate levels control root system architecture (Boer et al., 2020, Crawford and Glass, 1998). When nitrate levels are low, plants forage for nitrogen with increased primary root lengths and increased lateral root numbers. In contrast, high nitrate levels result in reduced lateral root elongation. Nitrate seemingly regulates primary and lateral root growth through two major phytohormones, auxin and cytokinin (Abualia et al., 2023, Sakakibara, 2021). Several nitrate transporters and sensors play a critical role in modulating this root growth pattern in response to nitrate. For example, NRT1.1/CHL6, a well-known nitrate transporter can sense and transport nitrate in addition to active auxin, IAA (Krouk et al., 2010). Auxin promotes primary and lateral root development, whereas cytokinin restricts lateral root development (Abualia et al., 2023, Michniewicz et al., 2019). Cytokinin restricts lateral root branching by limiting the availability of the auxin precursor indole-3-butyric acid (Michniewicz et al., 2019, Damodaran and Strader, 2019) and by altering PIN transport of IAA (Abualia et al., 2022). The concerted activities of these two hormones appear to drive nitrate-regulated root system architecture.

In this work, we report that elevated nitrate promotes anchor root formation. We also find that cytokinin acts antagonistically to nitrate in anchor root production by decreasing IBA contributions to the pool of active auxin in the collet. Overall, these data suggest a mechanism by which elevated nitrate can activate production of a shallow root system originating from anchor roots.

## 2 Materials and Methods

### 2.1 Plant Materials

Arabidopsis lines were in the Columbia (Col-0) background, including *ech2 ibr10* (Strader et al., 2011), *tob1-1* (Michniewicz et al., 2019), pTOB1:YFP-TOB1 (Michniewicz et al., 2019), *tob1 ech2 ibr10 (Michniewicz et al., 2019), ahk3 ahk4* (Nishimura et al., 2004), *arr3,4,5,6,7,8,9,15* (Zhang et al., 2011), *35s::KMD2-13(KMD2ox)* (Kim et al., 2013), *KMD2ox* pTOB1:YFP-TOB1 (Damodaran and Strader, 2024), and *KMD2ox ech2 ibr10* (Damodaran and Strader, 2024). Columbia (Col-0) was used as the wild type (Wt) in all assays.

### 2.2 Plant Growth Conditions and Phenotypic Assays

Seeds were surface sterilized using 20% bleach for 10 min, followed by several rinses with sterile water (Last and Fink, 1988). Seeds were then re-suspended in 0.1% agar and stratified for 3 days at 4 °C prior to plating on plant nutrient (PN) media (Haughn and Somerville, 1986) supplemented with 0.5% sucrose (PNS). To modulate nitrate, we replaced the 2 mM Ca(NO_3_)_2_ in the PN media recipe with 2 mM CaCl_2_ and added the appropriate amounts of KNO_3_ for the nitrate concentrations indicated. Because nitrate-free PNS medium has sufficient potassium for Arabidopsis growth (2.5 mM), we did not supplement with additional potassium when depleting KNO_3_. All assays were performed under constant light conditions (100 µMol/m^2^/s) at 22 °C. For IBA and IAA treatments, the seedlings were grown under yellow long-pass filters at 22 °C to prevent the degradation of indolic compounds.

To assess the long-term effect of nitrate on anchor root formation, seedlings were first grown for 3 days on half-strength MS media prior to transfer to sterile cylinders containing PNS media at indicated nitrate concentration. Anchor roots were counted 21 days after germination.

For assessing the effect of changing the nitrate levels, the seedling grown in low (0 mM NO_3_) and high (20 mM NO_3_) nitrate were transferred 4 days post germination to the mentioned concentration of either low or high nitrate. Anchor root formation was determined 9 days post germination.

To determine the effect of root tip excision, seedlings were grown in PNS media containing indicated concentration of nitrate and then the root tip was excised 5 days after germination (Jia et al., 2019). Anchor roots were then counted 8 days after germination.

### 2.3 Vector Construction and Transformation

The *ECH2* (*At1g76150*) upstream regulatory region was amplified using promECH2-F (5’-GGGAATTCGCTGGTGGTTAGATATAGA) and promECH2-R (5’-GGGTTTAAACCTCCGATCAGGATTAGAGCTC) (Supplementary Table 2). The resultant 570-bp product, which spanned the region between the stop codon of the neighboring genes and the *ECH2* start codon, was cloned into the pCR4-TOPO vector (Life Technologies) to create *pCR4-ECH2prom*. The *IBR10* (*At4g14430*) upstream regulatory region was amplified using promIBR10-F (GGGAATTCCTCATTGTCTTGTTGGGAG) and promIBR10-R (GGCTCGAGGGTGGTGATCGGAGGAAGA). The resultant 1310-bp product, spanning the region between *IBR10* and it most upstream neighboring gene, was cloned into pCR4-TOPO (Life Technologies) to generate *pCR4-promIBR10*. The *pCR4-promECH2* clone was digested with *EcoR*I and *Pme*I and pCR4-promIBR10 was digested using *EcoR*I and *Xho*I. These fragments were ligated into the pMCS:YFP-GW (Michniewicz et al., 2015) vector, digested with the analogous enzymes, to generate *promECH2:YFP-GW* and *promIBR10:YFP-GW*. Full length *ECH2* and *IBR10* cDNA were recombined into the pENTR-D-TOPO vector (Life Technologies) to generate *pENTR-D-ECH2 (Strader et al., 2011)* and *pENTR-D-IBR10*. These cDNA entry clones were used for LR reactions (LR Clonase, Life Technologies) into promECH2:YFP-GW and promIBR10:YFP-GW to create *promECH2:YFP-ECH2* and *promIBR10:YFP-IBR10*, which express N-terminal fusions of ECH2 and IBR10 driven behind their respective upstream regulatory regions. Recombinant plasmids were transformed into *Agrobacterium tumifaciens* strain *GV3101* (Koncz and Schell, 1986) through electroporation. Wild type *Arabidopsis thaliana* (Col-0) was transformed with these constructs using the floral dip method.(Clough and Bent, 1998) Transformants were selected in the presence of 10 µg/mL Basta (phosphinothricin) and lines homozygous with a single insert of the transgene were identified in subsequent generations. Individuals in the T_4_ generation were used for imaging.

### 2.4 Microscopy

Seedlings were mounted in water for imaging through a 20x/0.40 NA DRY objective on a Leica Thunder DMi8 inverted imaging system (Leica Microsystems, Germany) using LAS X software. The images were acquired in tile scanning mode using Leica-DFC9000GTC camera. The image acquisition settings including exposure time were kept constant across samples within each experiment. Post acquisition images were merged using LAS X software with the statistical merging method for further image processing in ImageJ software.

For confocal imaging seedlings were mounted in water for imaging through a 20x/0.75 NA dry objective on a Leica TCS SP8 confocal laser scanning microscope (Leica Microsystems, Germany) using LAS X (3.5.7.23225) software. The YFP fluorophores were excited using the 514 nm laser and emission was collected between 519 nm to 573 nm using HyD detectors. Images were acquired at 1024 x 1024 scan format and z-step intervals of 80 µm. The laser power and all acquisition settings were kept constant across samples within each experiment.

For peroxisomal co-localization, four-day-old seedlings expressing YFP-ECH2 or YFP-IBR10 were counterstained with 5 µM 8-(4-nitrophenyl)-BODIPY and excess dye was removed by rinsing in sterile water before mounting on a slide as described (Strader et al., 2011). The co-localization images were acquired using the 514 nm excitation laser for YFP and 488 nm excitation laser for BODIPY, with emission collected between 556 nm and 573 nm, and 494 nm and 516 nm respectively. To minimize spectral overlap the images were acquired using sequential scanning mode. The confocal images were merged, and scale bar was added using ImageJ software (Schneider et al., 2012).

### 2.5 Quantification and Statistical Analysis

The figure legends provide details of statistical analyses. Microsoft Excel and R (2024.04.0, “Chocolate Cosmos”) were used to perform statistical analyses for phenotypic assays. Graphs were generated using R and Microsoft Excel.

## 3 Results

### 3.1 Nitrate Regulates Anchor Root Formation

Given their importance in generating surface roots, we hypothesized that formation of anchor roots might be regulated by nitrogen status. As a first step, we evaluated whether seedling nitrate root responses under our lab conditions matched those previously published. In alignment with previous studies (Krouk et al., 2010, Boer et al., 2020, Giehl and von Wirén, 2014), we found that Arabidopsis seedlings displayed short root lengths under depleted nitrogen conditions (Figures 1A and 1B). At very low nitrate concentrations (0.01 to 0.5 mM), primary root lengths were longer than at elevated nitrate levels (Figures 1A and 1B). Similar to previous reports, (Gruber et al., 2013, Krouk et al., 2010) we observed increased numbers of lateral roots between 0.01 and 0.5 mM nitrate (Figure 1C), suggesting a foraging response. Based on these results, we were confident that our media and growth setup produced comparable developmental responses to previous studies.

**Figure 1.**
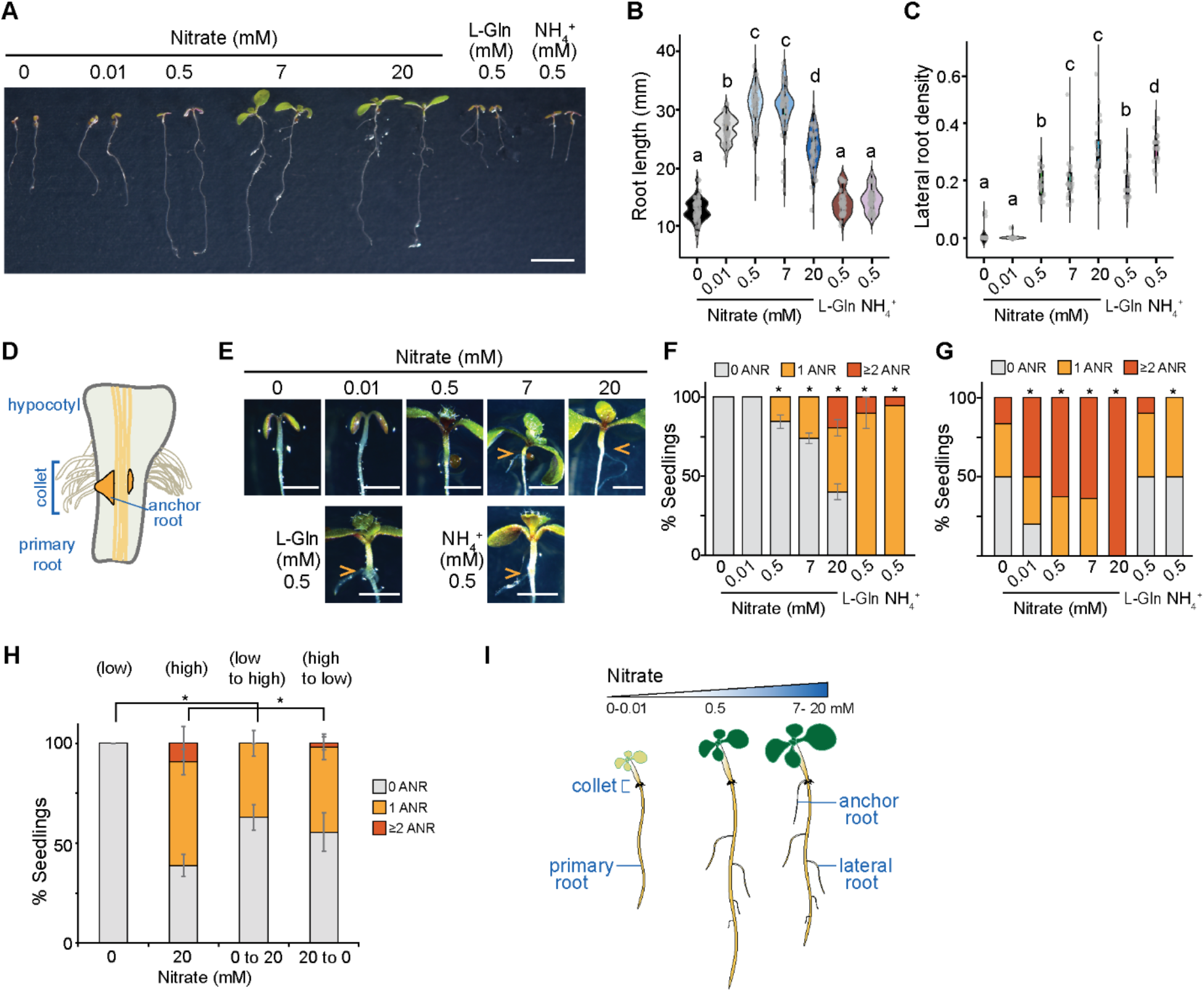
Nitrate regulates anchor root formation. (A) Photograph of wild type (Col-0) seedlings 8 days after germination on the indicated concentration of nitrate, ammonium, or glutamine. Scale bar = 1 cm. (B) Nitrate levels affect primary root growth of wild type (Col-0) seedlings. The violin plot combined with the box plot showed the primary root length of wild type (Col-0) seedlings 8 days after germination at the indicated concentrations of nitrate, ammonium, and glutamine. Statistical significance between the groups was determined by one-way ANOVA; each pair was compared using a Tukey Kramer HSD test and different letters indicate significant differences. Boxes show the first and third quartiles split by mean, and whiskers show the range. *n* ≥ 58 seedlings in total across three biological replicates and individual data points are shown as jitters. (C) Nitrate levels affect the lateral root density of wild type (Col-0) seedlings. The violin plot combined with the box plot showed the lateral root density (number of emerged lateral roots/cm of primary root length) of WT (Col-0) seedlings 8 days after germination at the indicated concentrations of nitrate, ammonium, and glutamine. Statistical significance between the groups was determined by one-way ANOVA; each pair was compared using a Tukey Kramer HSD test and different letters indicate significant differences. Boxes show the first and third quartiles split by mean, and whiskers show the range. *n* ≥ 58 seedlings in total across three biological replicates and individual data points are shown as jitters. (D) Graphical representation of an Arabidopsis collet with an emerged anchor root and anchor root primordia. (E) Close-up images of the collet of wild type (Col-0) seedlings 8 days after germination on the indicated concentration of nitrate, ammonium, or glutamine. Arrowheads indicate emerged anchor roots. Scale bars = 0.5 cm. (F) High nitrate level promotes anchor root development in wild type (Col-0) seedlings 8 days after germination. Quantification of number of anchor roots as percentage of seedlings with 0, 1, or ≥2 anchor roots. *n* ≥ 57 seedlings in total across three biological replicates and stacked bar graph represent the mean ± SD. Statistical significance between nitrate treatments was determined by comparing with 0 mM N treatment by chi-square test (*p ≤ 0.05). (G) Nitrate levels continuously alter anchor root development in 21-day-old wild type (Col-0). Quantification of number of anchor roots as the percentage of seedlings with either 0 or 1 or ≥2 anchor roots. *n* ≥ 11. (H) High nitrate promotes anchor root formation. Quantification of number of anchor roots as a percentage of seedlings with 0, 1 or ≥2 anchor roots. Seedlings were grown initially for 4 days in either 0 (low) or 20 mM NO_3_^−^ (high) and transferred to either low or high NO_3_^−^ PNS media. Anchor roots were counted 9 days after germination with n ≥ 54 seedlings in total across three biological replicates and stacked bar graph represents the mean ± SD. Statistical significance between different nitrate treatments was determined by chi-square tests (*p ≤ 0.05). (I) Graphical representation of Arabidopsis seedlings grown under varying nitrate levels affecting primary, lateral and anchor root development.

Because anchor roots have the potential to drive shallow exploration by root systems, and because nitrogen sources are differentially mobile, we questioned whether anchor root development would correlate with nitrate levels. Anchor roots emerge from the collet; we hypothesize that nitrate levels could alter this developmental program (Figure 1D). In wild type, anchor roots were rarely observed in seedlings grown under deficient and low nitrate conditions (0-0.01 mM NO_3_^−^; Figures 1E and 1F). Under optimal nitrate concentrations (2-7 mM NO_3_^−^), ∼15-20% of wild type seedlings developed anchor roots (Figures 1A, 1E and 1F). Under supraoptimal nitrate (i.e., 20 mM NO_3_^−^), most seedlings displayed increased anchor root formation (Figure 1F), even when sucrose was omitted from the media (Figure S1I). In addition, all seedlings displayed anchor roots when glutamine or ammonium were supplied (Figures 1F and S1A-D), with this effect appearing prominently at 21 days (Figure 1G). The persistence of nitrate effects on anchor root numbers at both 8 days and 21 days, combined with the fact that 7 mM NO_3_^−^ is sufficient for proper growth and yet have fewer anchor roots than seedlings grown in the presence of 20 mM NO_3_^−^ suggests that the differences observed are unlikely to be due solely to developmental delays at different nitrate levels. In addition, glutamine-induced anchor root formation was reduced with increasing nitrate levels, suggesting a complex interaction between nitrate and glutamine in controlling anchor roots (Figures S1E - S1H). Thus, elevated nitrate and/or the presence of alternate nitrogen sources promote anchor root formation concomitant with decreased primary root elongation.

Similar to superoptimal nitrate conditions, primary roots were short under nitrogen-free conditions (Figures 1A and 1B); however, there were no coincident anchor roots (Figures 1E–1F), suggesting that short roots alone are insufficient to stimulate anchor root formation. Further, we observed increased lateral root formation (Figure 1C) at nitrate concentrations that failed to stimulate anchor root development. When we transferred seedlings grown for four days in the absence of nitrate (0 mM N) to high nitrate (20 mM N), anchor root formation was promoted within 5 days in ∼70% of seedlings (Figure 1H). Although anchor roots and lateral roots are frequently considered analogous because they are both types of secondary roots, our data suggests that, unlike lateral roots, anchor root formation may not be a mechanism employed by root systems to scavenge nitrate (Figure 1I). Additionally, the fact that distinct conditions stimulate anchor and lateral root formation suggests that a combination of overlapping and distinct molecular mechanisms control development of these different secondary roots. The association between nitrate levels and anchor root formation encouraged us to pursue a deeper understanding of the molecular mechanism involved in nitrate-dependent anchor root development.

### 3.2 Nitrate Regulates Anchor Root Formation Through Cytokinin

Nitrate effects on root architecture are routed through cytokinin biosynthesis (Sakakibara, 2021, Abualia et al., 2023). Whereas cytokinin has a well-described role in restricting lateral root formation (Nenadić and Vermeer, 2021, Jing and Strader, 2019, Chang et al., 2015, Chang et al., 2013) and adventitious root formation (Lakehal et al., 2020, Damodaran and Strader, 2024), its role in anchor root formation had not previously been explored. Given the relationship between nitrate, cytokinin, and root system architecture, we decided to determine whether cytokinin signaling similarly mediated the effects of this nutrient on anchor root production to result in a shallow root system.

We first examined expression of the cytokinin response reporter *TCSn:GFP* and observed increased reporter signal in the collet under elevated nitrate conditions (Figure S2A and S2B), suggesting that cytokinin signaling in this region is regulated by nitrate levels. Cytokinin is perceived by the receptors ARABIDOPSIS HISTIDINE KINASE3 (AHK3) and AHK4, followed by a signaling cascade relayed through AHP (ARABIDOPSIS HISTIDINE PHOSPHOTRANSFER PROTEIN) which activates Type-B ARR (ARABIDOPSIS RESPONSE REGULATOR) transcriptional factors. The Type-A ARRs are negative regulators of cytokinin signaling (Figure 2A) (Kieber and Schaller, 2018). To determine cytokinin roles in anchor root development, we examined *ahk3 ahk4* and Type-A *arr* higher order mutants (Figure 2B and 2C), which are resistant and hypersensitive, respectively, to cytokinin (Nishimura et al., 2004, Zhang et al., 2011). In addition, we examined a line overexpressing *KISS ME DEADLY 2* (*KMD2*), which promotes TYPE-B ARR degradation and confers cytokinin resistance (Figure 2A) (Kim et al., 2013). We found that *ahk3 ahk4* and *KMD2ox* (*35S:KMD2-13*) developed anchor roots at lower nitrate concentrations than wild type (Figures 2B and 2C). In contrast, the cytokinin hypersensitive mutant *arr3,4,5,6,7,8,9,15* exhibited reduced anchor root formation at all examined nitrate levels (Figures 2B and 2C). Thus, like its effects on other secondary root systems, cytokinin restricts anchor root formation. Together these data suggest that cytokinin signaling appears to alter anchor root formation under varying nitrate concentrations.

**Figure 2.**
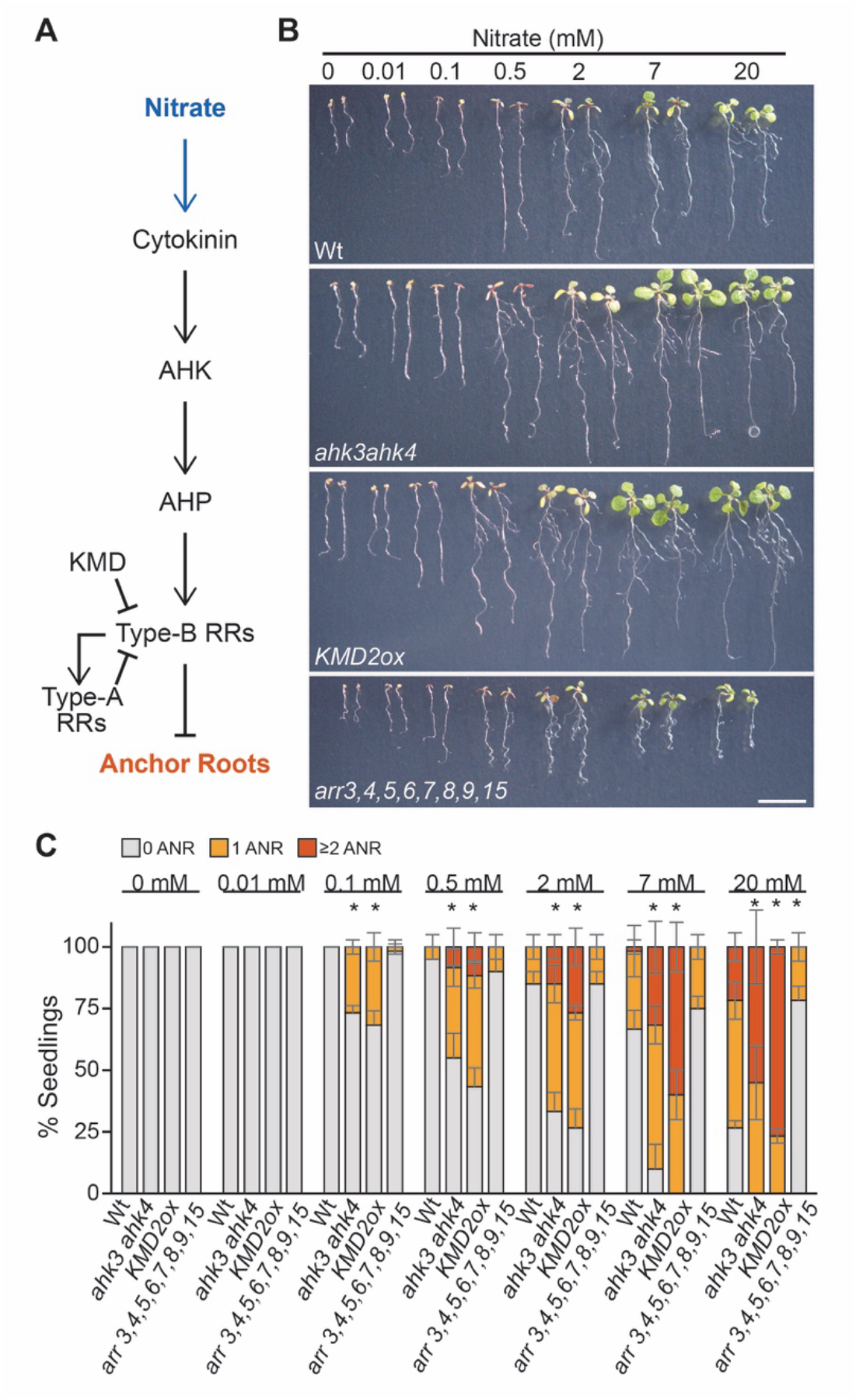
Cytokinin regulates anchor root development. (A) Representation of the cytokinin signaling pathway potentially involved in Arabidopsis anchor root development. (B) Photograph of 8-day-old seedlings of wild type (Col-0), *ahk3 ahk4*, *arr3,4,5,6,7,8,9,15*, or *KMD2ox* seedlings grown in the indicated concentration of nitrate. Scale bars = 1 cm. (C) Quantification of number of anchor roots 8 days post-germination for wild type (Col-0), *ahk3 ahk4*, *arr3,4,5,6,7,8,9,15*, or *KMD2ox*. The data represents the percentage of seedlings with 0, 1, or ≥2 anchor roots. *n* ≥ 60 seedlings in total across three biological replicates; stacked bar graph represents the mean ±SD. Statistical significance between genotypes under different nitrate treatments was determined by chi-square tests (*p ≤ 0.05).

### 3.3 Nitrate Regulates Anchor Root Formation Through TOB1/NPF5.12

TRANSPORTER OF IBA1 (TOB1/NPF5.12) acts downstream of cytokinin in both lateral root (Michniewicz et al., 2019) and adventitious root (Damodaran and Strader, 2024) development. TOB1/NPF5.12 transports both nitrate and the auxin precursor indole-3-butyric acid (IBA) and sequesters IBA in the vacuole to limit its availability for conversion to the active hormone auxin (Michniewicz et al., 2019). Loss of *TOB1* results in increased lateral roots (Michniewicz et al., 2019), a more branched root system architecture (Michniewicz et al., 2019), and increased adventitious rooting (Damodaran and Strader, 2024). Because cytokinin is clearly important for directing nitrate-regulated root architecture and TOB1/NPF5.12 acts downstream of cytokinin in other contexts, we hypothesized that TOB1/NPF5.12 acts downstream of cytokinin in nitrate-regulated anchor root development.

We first examined the effect of nitrate on *tob1* root elongation and found that *tob1* mutants displayed longer roots than wild type from 0 mM to 2 mM NO_3_^−^ (Figure 3A and 3B), suggesting that TOB1 restricts primary root growth at low to sub-optimal nitrate. We previously found that *tob1* displays increased lateral roots at 7 mM NO_3_^−^, which is the concentration of nitrate present in the plant growth media used in our group (Michniewicz et al., 2019). To determine the role of TOB1/NPF5.12 in nitrate-regulated lateral root development, we examined the effect of nitrate levels on emerged lateral roots in the *tob1* mutant. Compared to wild type, *tob1* displayed increased lateral roots between 0.5 mM NO_3_^−^ and 7 mM NO_3_^−^ (Figure 3C), suggesting TOB1 restricts lateral root formation under foraging and optimal nitrate conditions.

**Figure 3.**
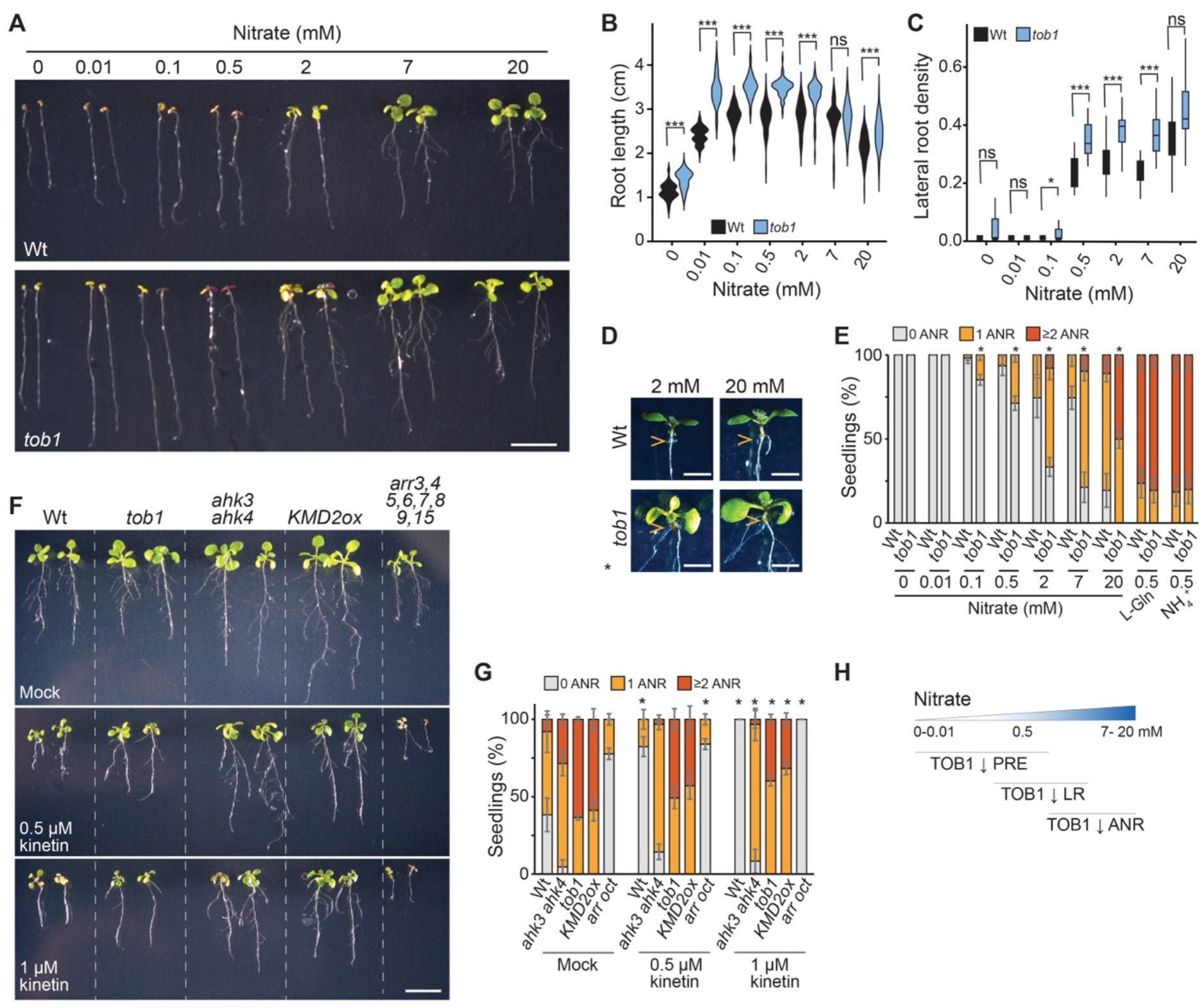
TOB1/NPF5.12 controls anchor root development under nitrate conditions through IBA availability. (A) Photograph of wild type (Col-0) and *tob1* seedlings 8 days after germination at the indicated concentrations of nitrate. Scale bar =1 cm. (B) Nitrate conditions affect primary root growth through TOB1/NPF5.12. The violin plot showed the primary root length of wild type (Col-0) and *tob1* seedlings 8 days after germination at the indicated concentrations of nitrate. Statistical significance between the groups was determined by one-way ANOVA, and significance between each pair was determined using the Wilcoxon signed-rank test (***P ≤ 0.001, ** P≤ 0.01, *P ≤ 0.05). (C) Nitrate conditions control lateral root density through TOB1/NPF5.12. Box plot shows the lateral root density (number of lateral roots per cm of primary root length) of wild type (Col-0) and *tob1* seedlings 8 days after germination at the indicated concentrations of nitrate. Statistical significance between the groups was determined by one-way ANOVA, and significance between each pair was determined using the Wilcoxon signed-rank test (***P ≤ 0.001, ** P≤ 0.01, *P ≤ 0.05). Boxes show the first and third quartiles split by mean, and whiskers show the range. (D) Close-up collet images of wild type (Col-0) and *tob1* seedlings 8 days after germination on the indicated concentration of nitrate. Arrowheads indicate the anchor root emerged from root-shoot junction/collet. (E) Quantification of number of anchor roots in wild type (Col-0) and *tob1* under different nitrate as a percentage of seedlings with 0, 1, or ≥2 anchor roots. *n* = 60 seedlings in total across three biological replicates and stacked bar graph represents the mean ±SD. Statistical significance between genotypes under different nitrate treatments was determined by chi-square tests (*p ≤ 0.05). (F) Resistance to cytokinin inhibition of anchor root by *tob1*. Photograph of wild type (Col-0), *tob1, ahk3 ahk4*, *KMD2ox* and *arr2,3,4,5,6,7,8,15* plants grown in the presence of 20 mM nitrate and the indicated concentration of kinetin. Scale bar = 1 cm. (G) Quantification of number of anchor roots in wild type (Col-0), *tob1*, *ahk3 ahk4*, *KMD2ox* and *arr2,3,4,5,6,7,8,15* as a percentage of seedlings with 0, 1, or ≥2 anchor roots, grown in 20 mM nitrate and treated at indicated concentration of kinetin. *n* = 52 seedlings in total across three biological replicates and stacked bar graph represents the mean ±SD. Statistical significance between genotypes and compared to mock treatment was determined by chi-square tests (*p ≤ 0.05). (H) Graphical representation of TOB1/NPF5.12 roles in the nitrate regulation of primary, lateral and anchor root development.

To determine the role of TOB1/NPF5.12 in nitrate-regulated anchor root formation, we examined wild type and *tob1* under varying nitrate conditions. We observed similar numbers of anchor roots in *tob1* as wild type at low nitrate levels, but elevated numbers of anchor roots compared to wild type at nitrate concentration ≥2 mM (Figures 3D and 3E). Overall, our data suggests TOB1/NPF5.12 regulates anchor root formation at optimal to high nitrate levels.

To determine whether cytokinin restricts anchor root formation through TOB1/NPF5.12 activity, we investigated the effect of exogenous cytokinin on anchor root formation in the *tob1* mutant. Cytokinin inhibits anchor root formation under high nitrate levels (20 mM NO_3_^−^) in wild type (Figures 3F and 3G). We found that similar to *ahk3 ahk4* and *KMD2ox*, *tob1* was resistant to the inhibitory effect of cytokinin on anchor root formation (Figure 3F and 3G). Thus, TOB1/NPF5.12 is necessary for the full inhibitory effects of cytokinin on anchor root formation, consistent with TOB1/NPF5.12 acting downstream of cytokinin in nitrate-regulated anchor root formation. Further, our data suggest that TOB1/NPF5.12 affects distinct aspects of root development at different nitrate concentrations (Figure 3H).

### 3.4 Nitrate Promotes *TOB1/NPF5.12* Expression in the Collet

The collet acts as a developmental boundary between shoot and root tissues and is located at the soil surface. Because we found that TOB1/NPF5.12 regulates anchor root formation, we tested whether nitrate levels affect *TOB1/NPF5.12* expression, which we previously found to be under the control of cytokinin signaling (Damodaran and Strader, 2024, Michniewicz et al., 2019).

Spatially-controlled *TOB1/NPF5.12* expression plays roles in both lateral root (Michniewicz et al., 2019) and adventitious root initiation (Damodaran and Strader, 2024). To understand whether nitrate levels regulate *TOB1/NPF5.12* expression in the collet, we examined a TOB1 translational reporter under different nitrate conditions. We observed that the YFP-TOB1 signal in the collet and collet root hairs strikingly increased with increasing nitrate concentrations (Figures 4A-C, S3A and S3B). This YFP-TOB1 signal primarily was present in collet root hair vacuoles (Figures 4B-4D), consistent with previously described roles for TOB1/NPF5.12 in sequestering IBA in the vacuole (Damodaran and Strader, 2024, Michniewicz et al., 2019).

**Figure 4.**
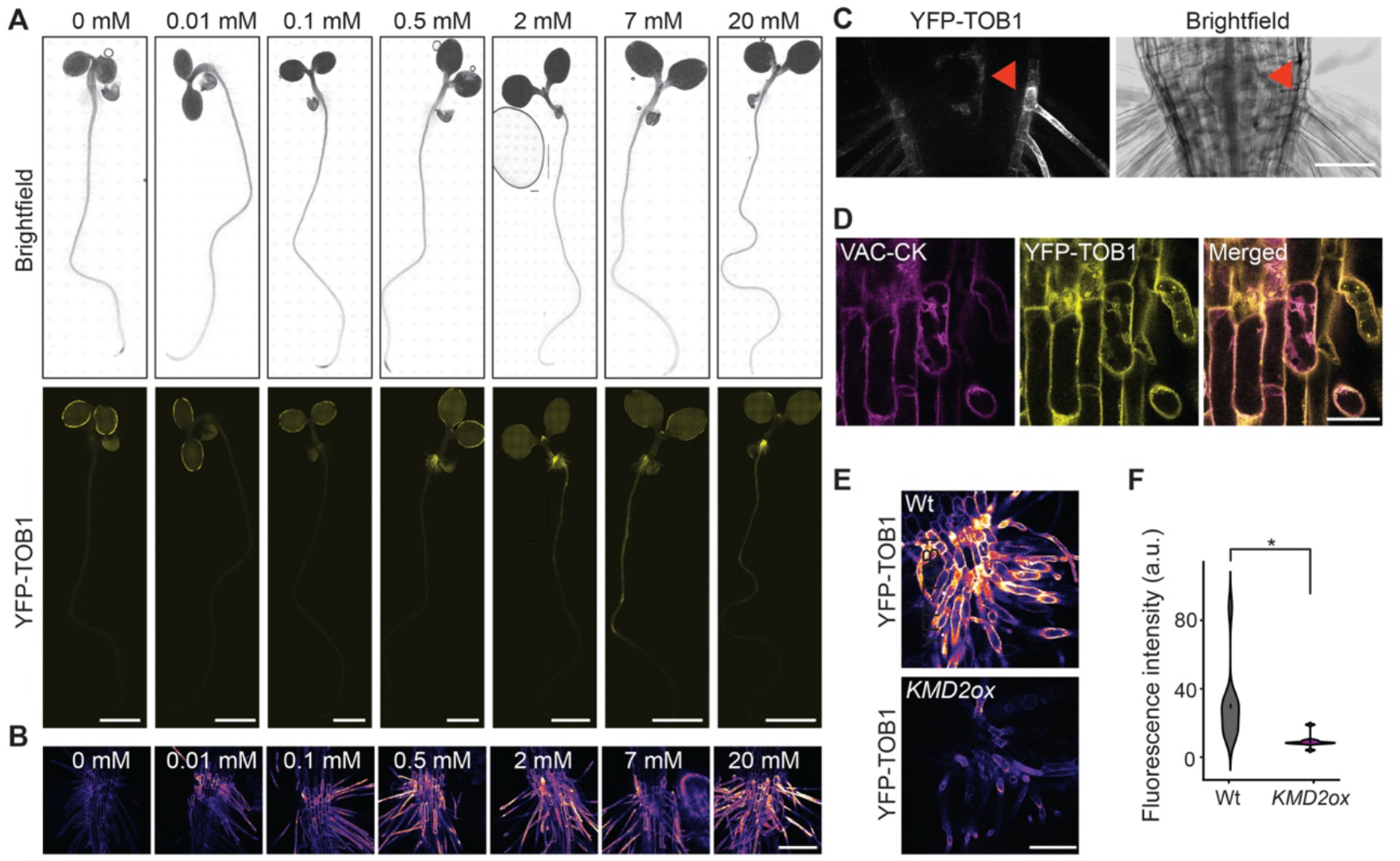
Nitrate promotes YFP-TOB1 expression in the collet. (A) Microscopy images of 4-day-old *tob1* seedlings expressing *pTOB1:YFP-TOB1* germinated on media containing the indicated nitrate concentrations. Scale bars = 0.25 cm. (B) TOB1/NPF5.12 expression in the collet under nitrate concentration. Confocal images of *pTOB1:YFP-TOB1* in the collet root hairs of 4-day-old seedling under different nitrate treatments. Scale bar = 100 µM. (C) TOB1/NPF5.12 expression in the collet root hairs and anchor root primordia. Confocal image of *pTOB1:YFP-TOB1* expression in the collet root hairs and anchor root primordia (indicated by arrowhead) of a 4-day-old seedling grown under 2 mM NO_3_^−^. Scale bar = 250 µM. (D) Colocalization of TOB1-YFP with a vacuolar marker. Confocal images of plants expressing *pTOB1:YFP-TOB1* and *VAC-CK* in the collet of 4-day-old seedling under 7 mM nitrate showing colocalization in the vacuolar membrane. Scale bar = 250 µM. (E) Reduced YFP-TOB1 signal in *KMD2ox*. Confocal images of *TOB1:YFP-TOB1* in 4-day-old wild type and *KMD2ox* lines grown in the presence of 20 mM NO_3_^−^. Scale bar = 250 µM. (F) Violin plot displaying the quantification of *TOB1:YFP-TOB1* fluorescence intensity in the collet of 4-day-old wild type and *KMD2ox* lines. n =10 seedlings; was a two-tailed t-test, *P <0.05.

Overall, these data are consistent with a model in which TOB1/NPF5.12 sequesters IBA at elevated nitrate levels to control auxin homeostasis in the collet.

*TOB1/NPF5.12* expression is regulated by cytokinin signaling in the context of lateral root (Michniewicz et al., 2019) and adventitious root development (Damodaran and Strader, 2024). We hypothesized that nitrate similarly controls anchor root development through cytokinin-regulated *TOB1* expression. We found that YFP-TOB1 signal was reduced in the collet of *KMD2ox* (Figure 4E and 4F), suggesting that nitrate-regulated *TOB1/NPF5.12* is dependent on cytokinin signaling.

Overall, our data suggest that TOB1/NPF5.12 acts downstream of nitrate and cytokinin signaling to connect soil nitrogen status to auxin homeostasis at the collet to regulate anchor root formation.

### 3.5 IBA-Derived Auxin Drives Anchor Root Formation

Auxin induces anchor root formation, consistent with previous findings that auxin signaling is critical to establishing emerging anchor roots (Jia et al., 2019). The auxin precursor IBA plays a critical role in lateral and adventitious root formation through its conversion to the active auxin indole-3-acetic acid (IAA) (Strader et al., 2010, Damodaran and Strader, 2024, Strader et al., 2011, Strader and Bartel, 2009). Our observation that TOB1/NPF5.12 activity regulates anchor root formation led us to hypothesize that IBA-derived auxin is critical for this process. To first identify IBA roles in anchor root formation, we observed the effect of exogenous IBA in anchor root formation. We grew wild-type seedlings in the presence of different concentrations of IAA and IBA; both the active auxin IAA and the auxin precursor IBA elicited anchor root formation (Figures 5A - 5C).

**Figure 5.**
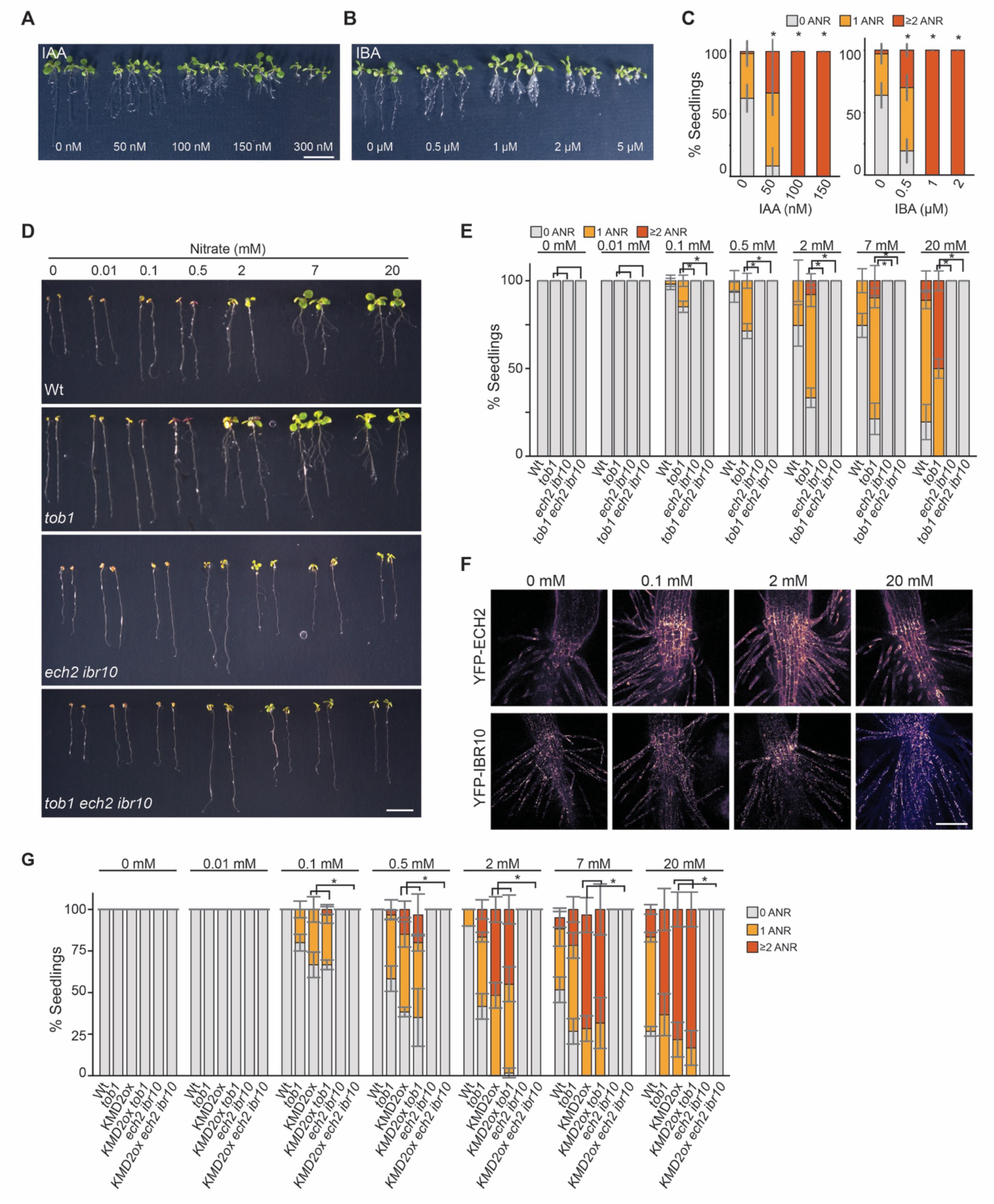
IBA-to-IAA conversion is essential for anchor root formation. (A) Photograph of wild type (Col-0) seedlings 8 days after germination in PNS media supplemented with the indicated concentrations of IAA and or a mock/ethanol treatment (0 µM). Scale bar = 1 cm. (B) Photograph of wild type (Col-0) plants 8 days after germination in PNS media substituted with indicated concentrations of IAA and mock/ethanol (0 µM). Scale bar, 1 cm. (C) Quantification of number of anchor roots in wild type seedlings grown in the presence of IAA and IBA as a percentage of seedlings with 0, 1, or ≥2 anchor roots. *n* = 52 seedlings in total across three biological replicates and data represent the mean ± SD. (D) Photographs of wild type (Col-0), *tob1*, *ech2 ibr10*, and *tob1 ech2 ibr10* plants grown for 8 days on media with indicated concentrations of nitrate. Scale bar = 1 cm. (E) Quantification of number of anchor roots in wild type (Col-0), *tob1*, *ech2 ibr10*, and *tob1 ech2 ibr10* as a percentage of seedlings with 0, 1, or ≥2 anchor roots. *n* = 54 seedlings in total across three biological replicates and data represent the mean ±SD. Statistical significance between genotypes and anchor root formation was determined by chi-square test (*p ≤ 0.05). (F) IBA-to-IAA conversion enzymes are expressed in the collet region. Confocal images of *pECH2:YFP-ECH2* and *pIBR10:YFP-IBR10* in the collet root hairs of 4-day-old seedlings grown in the presence of indicated concentration of nitrate. (G) Nitrate-cytokinin-IBA control anchor root formation. Quantification of number of anchor roots in wild type (Col-0), *tob1, KMD2ox, KMD2ox tob1, ech2 ibr10, KMD2ox ech2 ibr10* seedlings grown under different nitrate as the percentage of seedlings with 0, 1, or ≥2 anchor roots. *n* = 60 seedlings in total across three biological replicates and stacked bar graph represents the mean ±SD. Statistical significance between genotypes under different nitrate treatments was determined by chi-square tests (*p ≤ 0.05).

IBA is converted to IAA in a multistep enzymatic pathway including the peroxisomal enzymes ENOYL-CoA HYDRATASE2 (ECH2), INDOLE 3-BUTYRIC ACID RESPONSE1 (IBR1), IBR3, and IBR10 (Strader and Bartel, 2011). To determine whether endogenous IBA-derived IAA has a role in anchor root formation, we examined seedlings defective in the IBA-to-IAA conversion process. Although we observed that *ech2 ibr10* mutant plants could form a few lateral roots at 0.5 and 2 mM nitrate (Figure 5D and S4A), the mutant failed to display emerged anchor roots at any examined nitrate level (Figure 5E), suggesting that IBA-to-IAA conversion is critical for anchor root formation.

TOB1/NPF5.12 both sequesters IBA in the vacuole (Michniewicz et al., 2019) and restricts anchor root formation (Figure 3D). Because TOB1/NPF5.12 transports both IBA and nitrate (He et al., 2017, Michniewicz et al., 2019), we questioned whether TOB1/NPF5.12 roles in anchor root formation were due to its potential role in cellular nitrate homeostasis or in its role in regulating IBA contributions to the cellular auxin pool. To parse these two possibilities, we examined anchor roots of *tob1 ech2 ibr10* and found that a loss of IBA-to-IAA conversion suppressed the *tob1* anchor root phenotype (Figure 5D and 5E). Thus, IBA-to-IAA conversion acts downstream of TOB1 roles in anchor root regulation.

To explore the effect of nitrate on the spatial expression of IBA conversion enzymes, we observed ECH2 and IBR10 translational reporters at different nitrate levels. Both ECH2 and IBR10 have been proposed to perform the enoyl-CoA hydratase step of this conversion process and are likely redundant in this activity (Strader et al., 2011). We observed strong YFP-ECH2 signal in the collet, with the signal gradually diminishing towards the root tip (Figure S4B-S4D). Similarly, YFP-IBR10 signal was strongest in the collet (Figure S4E-S4G). Collet YFP-ECH2 signal decreased with increasing nitrate levels, whereas collet YFP-IBR10 signal was unaffected by the tested nitrate concentrations (Figure 5F, S4C, S4D, S4F and S4G). As previously reported, these reporters localized to the peroxisome (Figure S3H), which is the site of IBA-to-IAA conversion (Strader et al., 2011, Zolman et al., 2008). These expression data and the mutant phenotypes suggest local IBA-to-IAA conversion in the collet promotes anchor root formation.

Because TOB1/NPF5.12 acts downstream of cytokinin signaling and IBA-to-IAA conversion acts downstream of TOB1/NPF5.12 activity in regulating anchor root formation, we wanted to confirm that IBA-to-IAA conversion acts downstream of cytokinin in this process. We found that, blocking IBA-to-IAA conversion with the *ech2* and *ibr10* mutations suppressed *KMD2ox* nitrate-regulated anchor root phenotypes (Figure 5G and S4I). Consistent with this result, the *KMD2ox tob1* line resembled both *KMD2ox* and *tob1* parents (Figure 5G and S4I), suggesting that those two factors act in the same genetic pathway. We hypothesize that the more dramatic phenotype of *KMD2ox* could be caused by disruption of multiple targets of cytokinin signaling (of which *TOB1* is one) and/or by redundancy in the *TOB1* family (Michniewicz et al., 2019). Together, these data indicate that cytokinin signaling via KMD2 is upstream of IBA transport by TOB1/NPF5.12 and IBA-to-IAA conversion pathway to control anchor root formation at different nitrate levels.

Previously, root tip excision was shown to promote anchor root formation (Jia et al., 2019). We hypothesized that nitrate-induced anchor root formation might be distinct from excision-induced anchor root formation. Indeed, we found that excision and elevated nitrate had additive effects on anchor root formation (Figure S5), consistent with the possibility that these stimuli act independently to promote these collet-borne roots. Further, IBA-to-IAA conversion, which is deficient in the *ech2 ibr10* mutant, is required for both nitrate- and excision-induced anchor root formation (Figure S5).

### 3.6 Multiple Species Generate Nitrate-Regulated Anchor Roots

To better understand the effects of nitrate on anchor root formation in other plants, we examined the effects of nitrate on canola, which, like Arabidopsis, is in the *Brassicaceae* family, and tomato, which is in the *Solanaceae* family. We chose to examine these plants because they both generate large seeds and have less reliance on exogenous nitrogen sources to fuel early seedling growth. We found that canola seedlings produced anchor roots in the absence of exogenous nitrate and displayed nitrate-regulated increases in anchor root formation (Figure 6A and 6B). Likewise, tomato seedling formation of anchor roots is stimulated by elevated nitrate (Figure 6C and 6D), although this increase was accompanying by decreased primary root elongation. These data further suggest that nitrate-regulated anchor root formation is common to multiple dicots.

**Figure 6.**
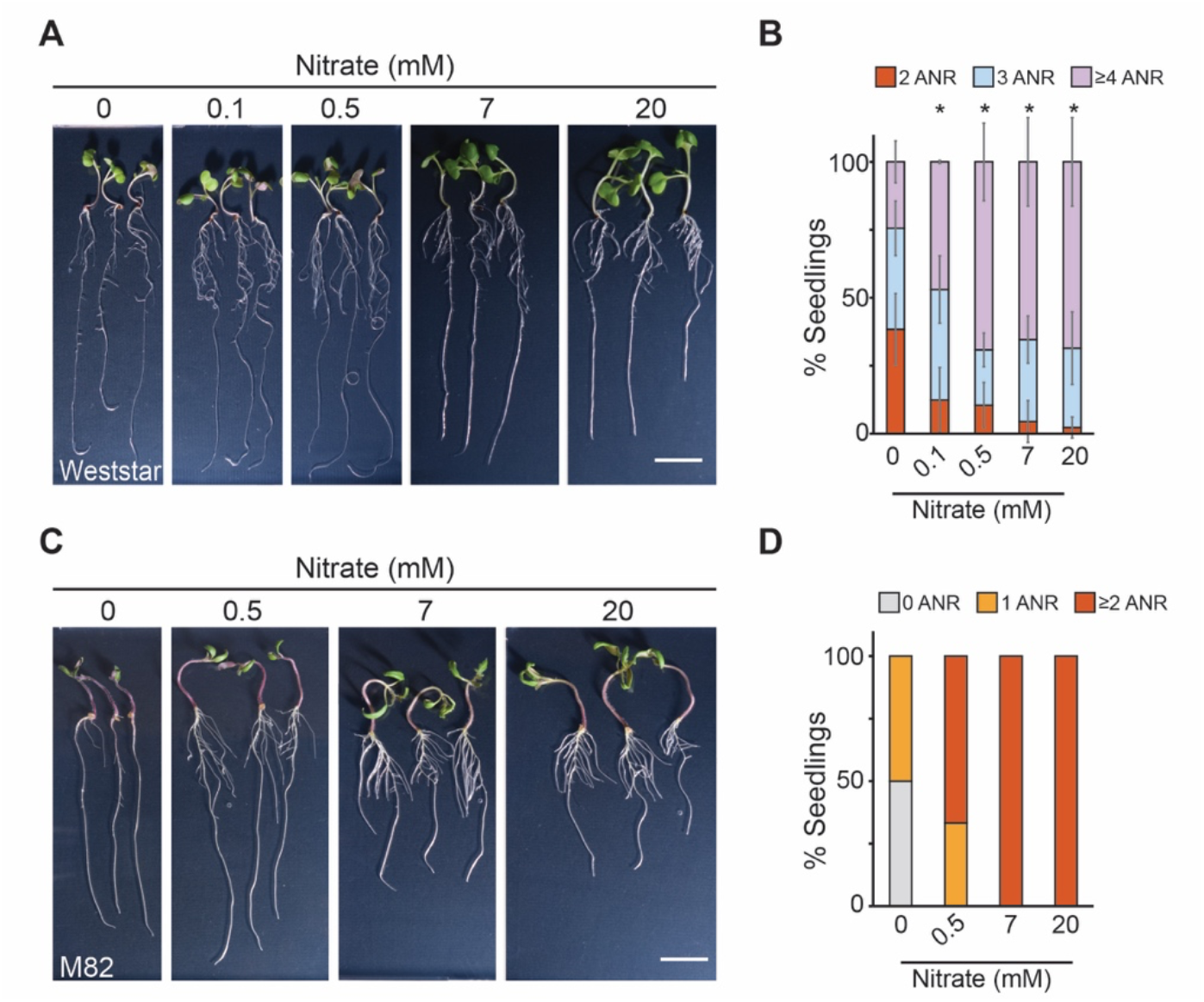
Nitrate regulates anchor roots in canola and tomato. (A) Photographs of 10-day-old canola seedlings grown in the presence of the indicated nitrate concentrations. Scale bar = 3 cm. (B) Quantification of anchor roots from 10-day-old canola seedlings grown in the presence of the indicated nitrate concentrations as the percentage of seedlings with 0, 1, or ≥2 anchor roots. *n* = 60; bars represent mean ±SD from three biological replicates. Statistical significance in anchor root formation under different nitrate treatment was determined by chi-square test (*p ≤ 0.05). (C) Photographs of 10-day-old tomato seedlings grown in the presence of the indicated nitrate concentrations. Scale bar = 3 cm. (D) Quantification of anchor roots from 10-day-old tomato seedlings grown in the presence of the indicated nitrate concentrations as the percentage of seedlings with 0, 1, or ≥2 anchor roots. *n* =12.

## 4 Discussion

Root system architecture (RSA) is highly plastic, which allows plants to optimize nutrient and water acquisition. Nitrate availability is critical regulator of RSA. Abundant nitrate near the soil surface results in a shallow RSA, whereas nitrate deficiency drives a deeper root system (Shahzad and Amtmann, 2017, Gruber et al., 2013). Low nitrate promotes a foraging response, leading to primary and lateral root elongation whereas elevated nitrate reduces primary and lateral root elongation (Gruber et al., 2013, Giehl and von Wirén, 2014). Although previous studies have described how nitrate shapes primary and lateral root development, the role of anchor roots in this process has been largely overlooked.

In this work, we show that anchor roots, which emerge from the collet, are regulated by nitrate availability. Anchor root formation displays a marked increase under high nitrate or in the presence of alternate nitrogen sources like ammonium or glutamine. Thus, anchor roots might serve a complementary role in RSA remodeling when nitrogen is replete near the soil surface. This functional complementarity is analogous to root class partitioning described in cereals, where anatomically distinct root types occupy different soil horizons and are regulated by both developmental programs and environmental signals (Lynch, 2013).

Unlike lateral roots that arise from premarked pericycle cells exiting from the root meristem, anchor roots originate from the collet, which was established during embryogenesis. Whereas excision of the root tip promotes the anchor root emergence in Arabidopsis seedlings (Lucas et al., 2011, Jia et al., 2019), our work reveals that additional mechanisms contribute to anchor root emergence for nutrient uptake. In particular, we found that cytokinin is a negative regulator of anchor root formation under nitrate-replete conditions (Figure 2). Canonical cytokinin signaling through AHK3/4 receptors and downstream type-B ARRs restricts anchor root formation, placing cytokinin as an important modulator of anchor root formation. This result is consistent with emerging evidence that cytokinin acts as a context-dependent rheostat in root development decisions (Růžička et al., 2009, Ioio et al., 2008).

The NRT1/PTR family of transporters transport nitrate and also a broad range of substrates, including phytohormones (Leran et al., 2014). The NRT1.1/ CHL6 transceptor is a key transporter involved in nitrate perception and regulates root system architecture under varying nitrate conditions by controlling IAA transport (Krouk et al., 2010). We also identify another member of this large family, TOB1/NFP5.12, a transporter of both IBA and nitrate (Michniewicz et al., 2019), as an integrator of nitrate and cytokinin control of nitrate-regulated root branching.

TOB1/NPF5.12 is required both for lateral root responses at low-to-optimal nitrate and for proper repression of anchor roots at high nitrate (Figure 7B). *TOB1/NPF5.12* expression in the collet increases with nitrate and localizes to the vacuole of collet root hairs, suggesting a role in nitrate and auxin precursor homeostasis at the root-shoot junction. Loss of *TOB1/NPF5.12* confers cytokinin resistance and results in increased nitrate-regulated anchor root formation, placing TOB1/NPF5.12 downstream of cytokinin in this developmental pathway. The dual capacity of TOB1/NPF5.12 to transport both a macronutrient (nitrate) and a hormone precursor (IBA) places it within an emerging class of multifunctional transport proteins that may couple nutrient sensing to hormone homeostasis (Krapp, 2015). Its vacuolar localization in collet root hairs further suggests that IBA sequestration, rather than degradation or redistribution to other tissues, is the primary mechanism for limiting IBA-to-IAA conversion under high nitrate, providing a rapid and potentially reversible regulatory mode well-suited to the dynamic nature of soil nitrate availability. Aligned with the role of TOB1/NPF5.12 as an IBA transporter, we further uncover that IBA-to-IAA conversion is essential for anchor root development and that this conversion acts downstream of cytokinin signaling and TOB1/NPF5.12 activity. Mutants defective in the IBA conversion pathway are unable to efficiently form anchor roots even under high nitrate or after root tip excision, reinforcing the role of IBA metabolism as a checkpoint in anchor root development. Together these results suggest the complexity of nitrate and additional substrates transport regulated by NPF transporters in nitrate dependent root growth.

**Figure 7.**
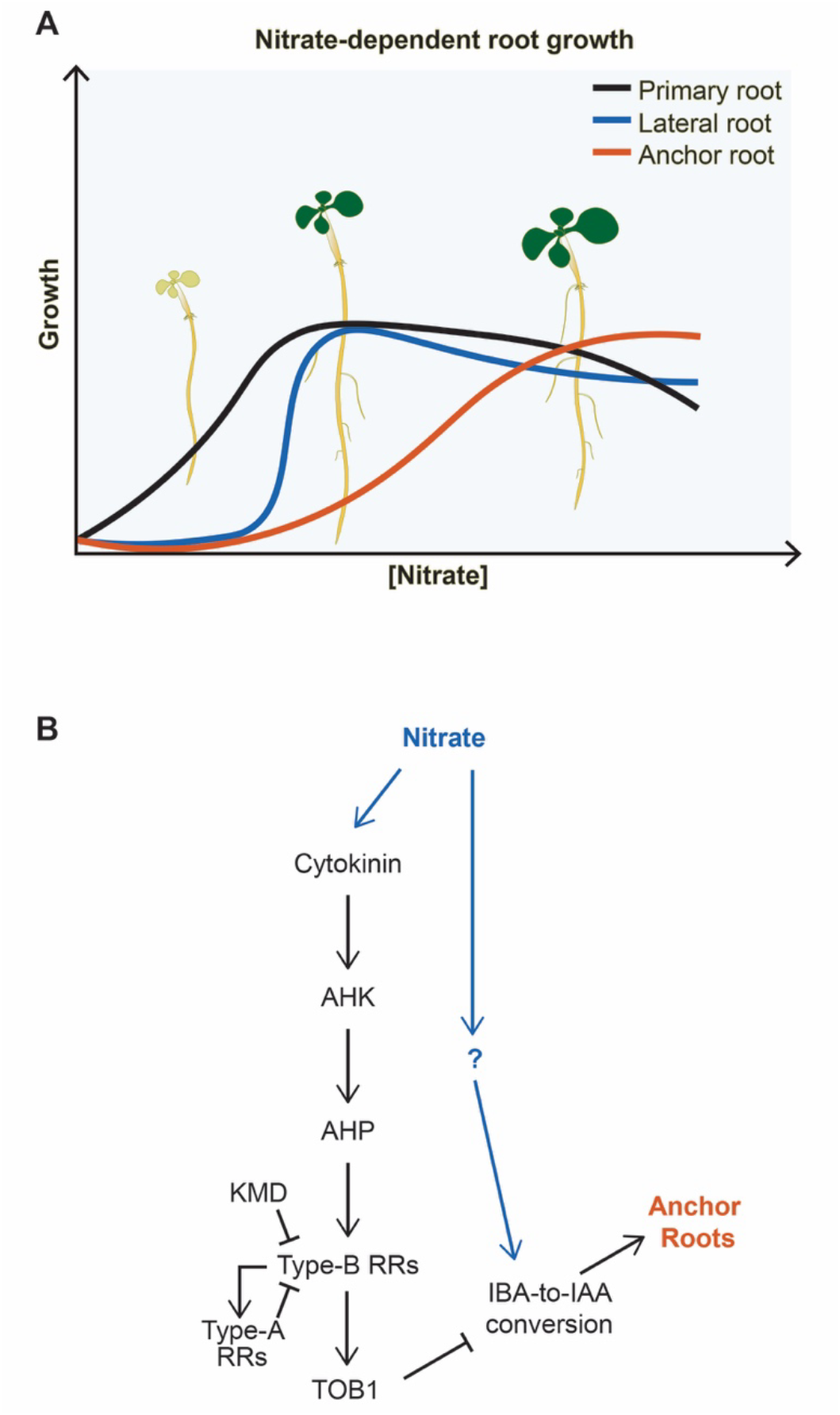
Models of nitrate effects on root elongation, lateral roots, and anchor roots. (A) Graphical representation of nitrate dependent root growth. Under nitrate-free conditions, root growth is inhibited. Under suboptimal nitrate, foraging conditions promote primary and lateral root growth. Supraoptimal nitrate results in reduced primary root elongation and lateral root production while promoting anchor root production. (B) Nitrate controls anchor root formation through IBA homeostasis. Graphical representation of the molecular mechanism governing nitrate-dependent anchor root emergence. Cytokinin restricts anchor root emergence by promoting TOB1/NPF5.12 expression through its signaling pathway. TOB1/NPF5.12 reduces the contribution of IBA to active auxin, which is necessary for anchor root emergence at high nitrate levels. Defects in IBA-to-IAA conversion affect anchor root formation even under high nitrate levels, suggesting an additional unknown pathway regulating anchor roots.

This model adds to our previous understanding that elevated nitrate results in upregulated cytokinin biosynthesis; which is then transported to the shoot to promote polar auxin transport (Sakakibara, 2021). The resultant rootward auxin transport alters primary root elongation and lateral root formation in response to elevated nitrate conditions (Abualia et al., 2023, Sakakibara, 2021). In contrast to this distal signaling model of nitrate-regulated lateral root production, we hypothesize that nitrate-regulated anchor root production at the soil surface is driven by local cytokinin and auxin signaling.

We propose a model in which nitrate availability promotes anchor root formation, which requires IBA-derived auxin (Figure 7B). In parallel, a nitrate-regulated braking mechanism is activated through cytokinin signaling, which in turn promotes *TOB1* expression to sequester IBA in vacuoles, thereby limiting its conversion to active IAA. This suppresses anchor root formation and biases RSA. When this braking mechanism is deficient, as in the case of the *ahk3 ahk4* or *tob1* mutants, there is hyperproduction of nitrate-induced anchor roots. The enrichment of *ECH2* and *IBR10* expression in collet root hairs (Figures 5F, S4D and S4G) supports this spatial model and suggests that the collet region acts as a key integration hub for hormonal and nutrient cues. This local signaling model complements the systemic nitrate-cytokinin-auxin cascade described for lateral root regulation (Abualia et al., 2023). Together, they reveal that the same cue can drive distinct RSA outcomes by engaging both systemic and local hormone networks.

Nitrate is a highly mobile nutrient that leaches through the soil column. Thus, its acquisition is strongly time sensitive. When nitrate availability is low at the soil surface, it is likely that available nitrate has already moved deeper into the soil profile, favoring investment in deeper primary and lateral root growth. Conversely, when nitrate is abundant at the soil surface, rapid acquisition prior to leaching becomes advantageous. We therefore hypothesize there is a time-critical incentive for shallow rooting, making investment in anchor roots could be beneficial for capturing transient nitrate pulses.

A limitation of this study is that anchor root development has been examined in seedlings grown under continuously fixed nitrate levels in agar medium, which does not replicate the soil environment. Future experiments focused on translating these assays to soil or rhizotrons could help understand role of metabolites, growth stage and phytohormones with altering nutrients in modifying RSA through anchor roots.

Several important questions emerge from this work. The identity of the nitrate sensor(s) acting at the collet has not been established. Of particular translational importance is whether the regulatory logic described here is conserved in crop species. Comparative studies testing whether cytokinin signaling, IBA metabolism, and TOB1 homologs regulate these structures in crop species would advance the translational significance of this framework.

Together, these findings uncover a distinct developmental mechanism regulating anchor root formation at the collet region under varying nitrate levels, providing a framework for future studies on nutrient acquisition in complex environments. Understanding how distinct root types respond to environmental signals will be essential for engineering crops with tailored root systems for efficient nutrient capture and resilience in variable soils.

## Acknowledgements

We want to thank Joe Kieber for providing cytokinin mutant seeds and Suyash Patil and Lena Mueller for the tomato and canola seeds. We also thank Yiling Fang, Yuren Cao, Ryan Bitter, Eduardo Flores, Sunita Pathak, Jhonny Figueroa Parra, Se-Hwa Lee, David Korasick and Andrew Willoughby for critical comments on the manuscript. This research was supported by the National Science Foundation (PGRP BIO-2112056 to LCS), the National Institutes of Health (R35 GM136338 to LCS), by gifts to the Salk Institute’s Harnessing Plants Initiative (HPI) from the Bezos Earth Fund, the Hess Corporation, and the TED Audacious Project.

---

**Supplemental Figure 1.**
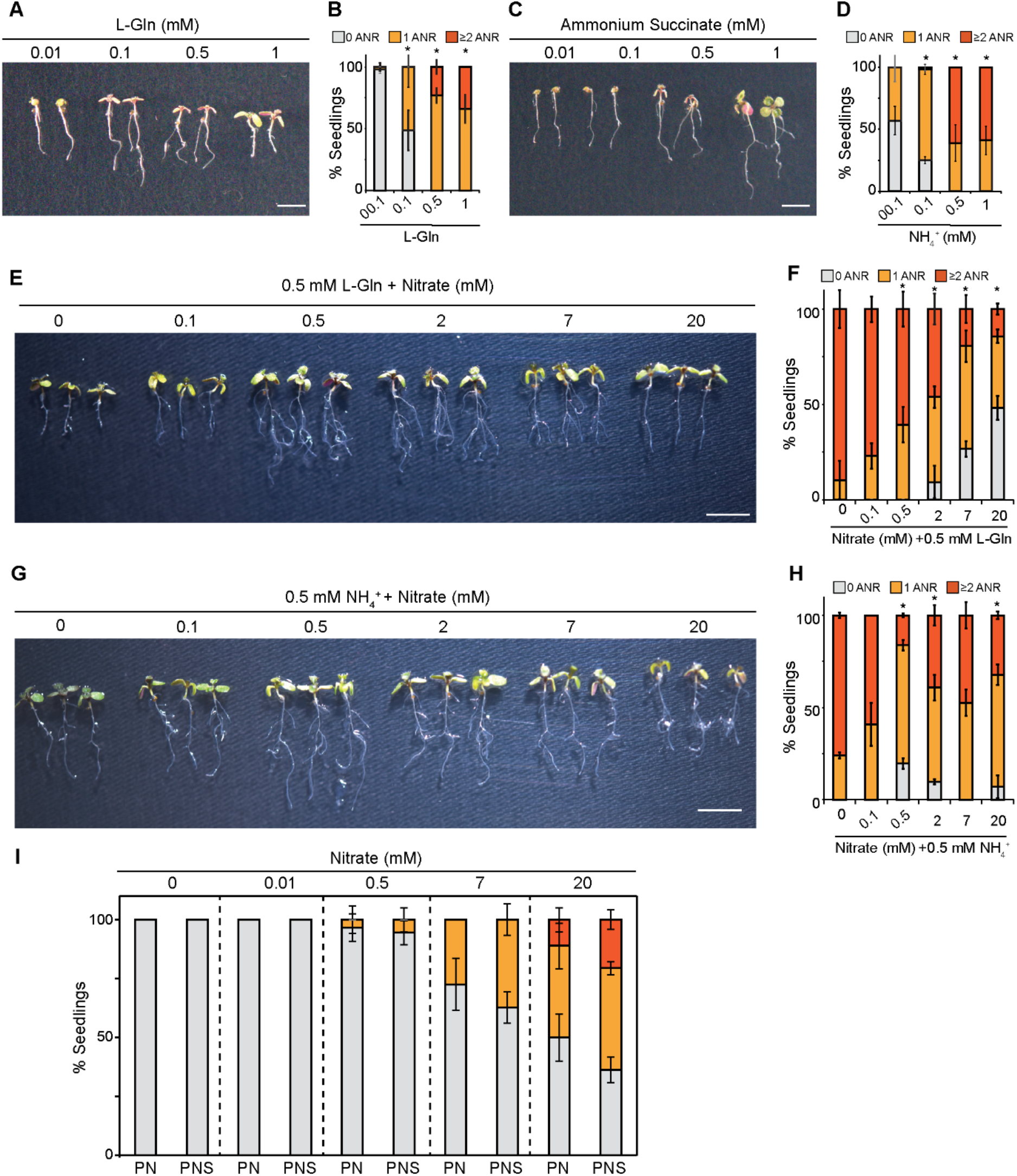
Organic nitrogen sources promote anchor root development. (A) Photograph of 8-day-old Arabidopsis (Col-0) seedlings grown in the presence of the indicated concentrations of L-Gln. Scale bar = 1 cm. (B) Quantification of number of anchor roots in wild type seedlings grown in different concentration of L-Gln shown as percentage of seedlings with 0, 1, or ≥2 anchor roots. *n* = 56 and data represents mean ±SD. Statistical significance in anchor root formation under different L-gln treatment was determined by chi-square test (*p ≤ 0.05). (C) Photograph of 8-day-old wild type seedlings grown in different concentration of ammonium succinate. Scale bar = 1 cm. (D) Quantification of number of anchor roots in wild type seedlings grown in different concentration of ammonium succinate shown as percentage of seedlings with 0, 1, or ≥2 anchor roots. *n* = 60 and data represents mean ±SD. Statistical significance in anchor root formation under different ammonium succinate treatment was determined by chi-square test (*p ≤ 0.05). (E) Photograph of 8-day-old wild type seedlings grown in the presence of 0.5 mM L-Gln and the indicated concentration of nitrate. Scale bar = 1 cm. (F) Quantification of number of anchor roots in wild type seedlings grown in 0.5 mM L-Gln and indicated concentration of nitrate shown as percentage of seedlings with 0, 1, or ≥2 anchor roots. *n* = 60 and data represents mean ±SD. Statistical significance in anchor root formation under different treatment was determined by chi-square test (*p ≤ 0.05). (G) Photograph of 8-day-old wild type seedlings grown in the presence of 0.5 mM ammonium succinate and the indicated concentration of nitrate. Scale bar = 1 cm. (H) Quantification of number of anchor roots in wild type seedlings grown in 0.5 mM ammonium succinate and indicated concentration of nitrate. *n* = 62 and data represents mean ±SD. Statistical significance in anchor root formation under different treatment was determined by chi-square test (*p ≤ 0.05). (I) Quantification of number of anchor roots in wild type seedlings grown in PN media (without sucrose) and PNS (with sucrose) with indicated concentration of nitrate. *n = 53* and data represents mean ±SD. Statistical significance in anchor root formation under different treatment was determined by chi-square test (*p ≤ 0.05).

**Supplemental Figure 2.**
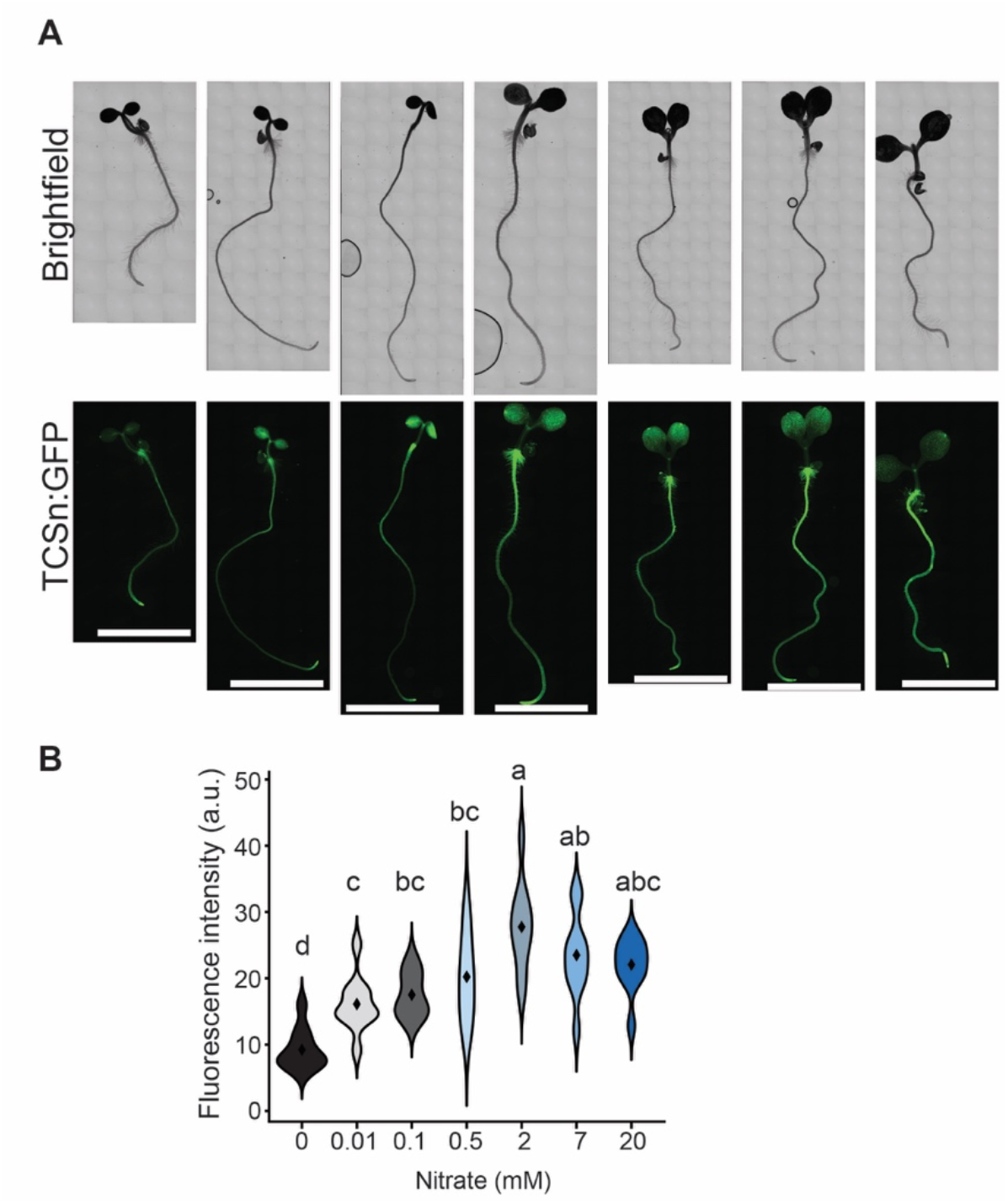
Nitrate promotes TCSn:GFP expression in the collet. (A) Microscopy images of 4-day-old seedlings expressing TCSn:GFP germinated on media containing the indicated nitrate concentrations. Scale bars = 0.5 cm. (B) Violin plot displaying quantification of fluorescence intensity in collet tissues of TCSn:GFP expressing 4-day-old seedlings under different nitrate concentrations. n =10 seedlings and statistical significance between the groups was determined by one-way ANOVA; each pair was compared using a Tukey Kramer HSD test and different letters indicate significant differences.

**Supplemental Figure 3.**
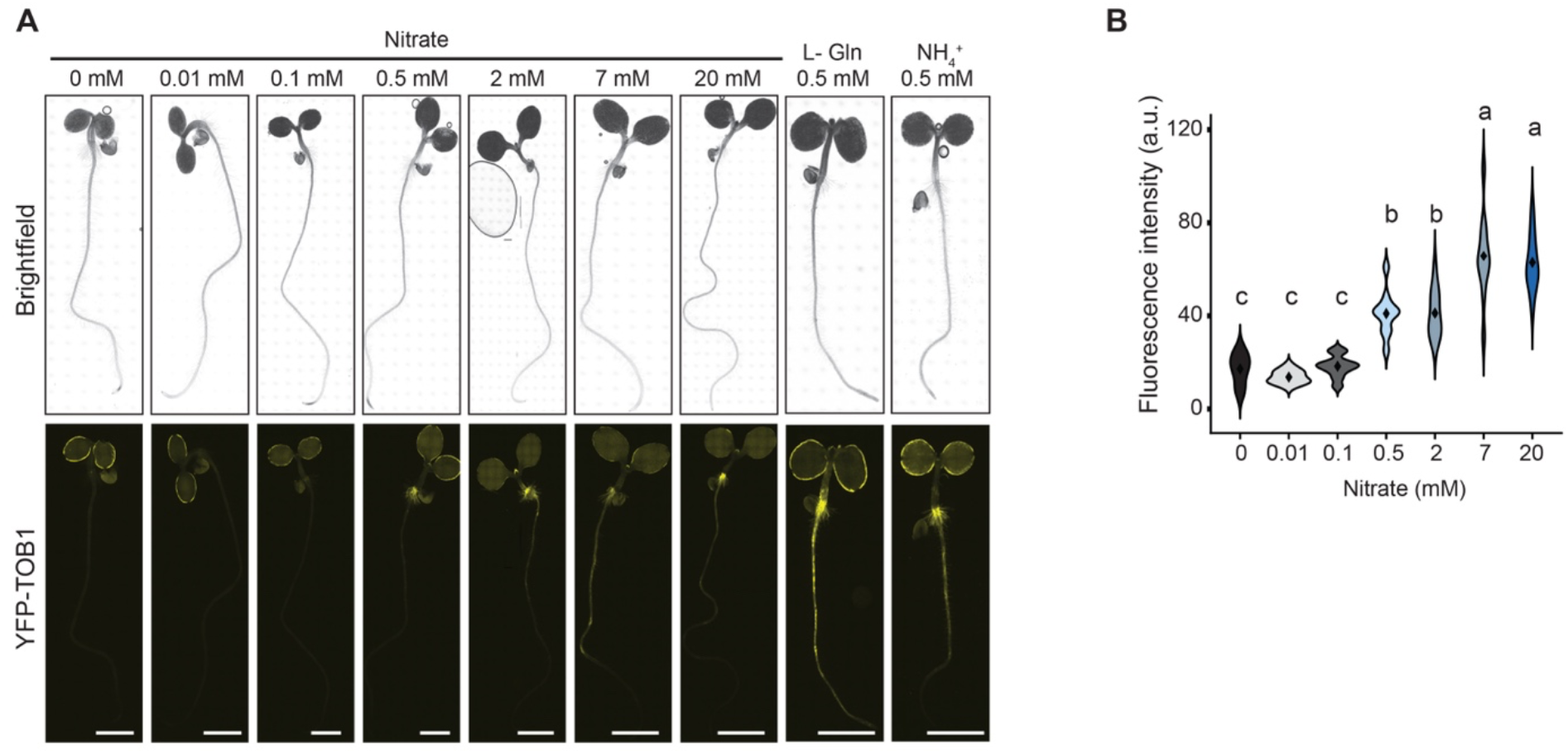
*TOB1/NPF5.12* expression is induced by nitrate and organic nitrogen. (A) Microscopy images of 4-day-old *pTOB1:YFP-TOB1*-expressing seedlings grown with the indicated concentrations of nitrate, L-Gln, or ammonium. Scale bars = 1 mm. (B) Violin plot displaying quantification of *TOB1:YFP-TOB1* fluorescence intensity in collets of 4-day-old seedlings grown under the indicated nitrate concentrations. *n* =10 seedlings; statistical significance between the groups was determined by one-way ANOVA; each pair was compared using a Tukey-Kramer HSD test and letters indicate significant differences.

**Supplemental Figure 4.**
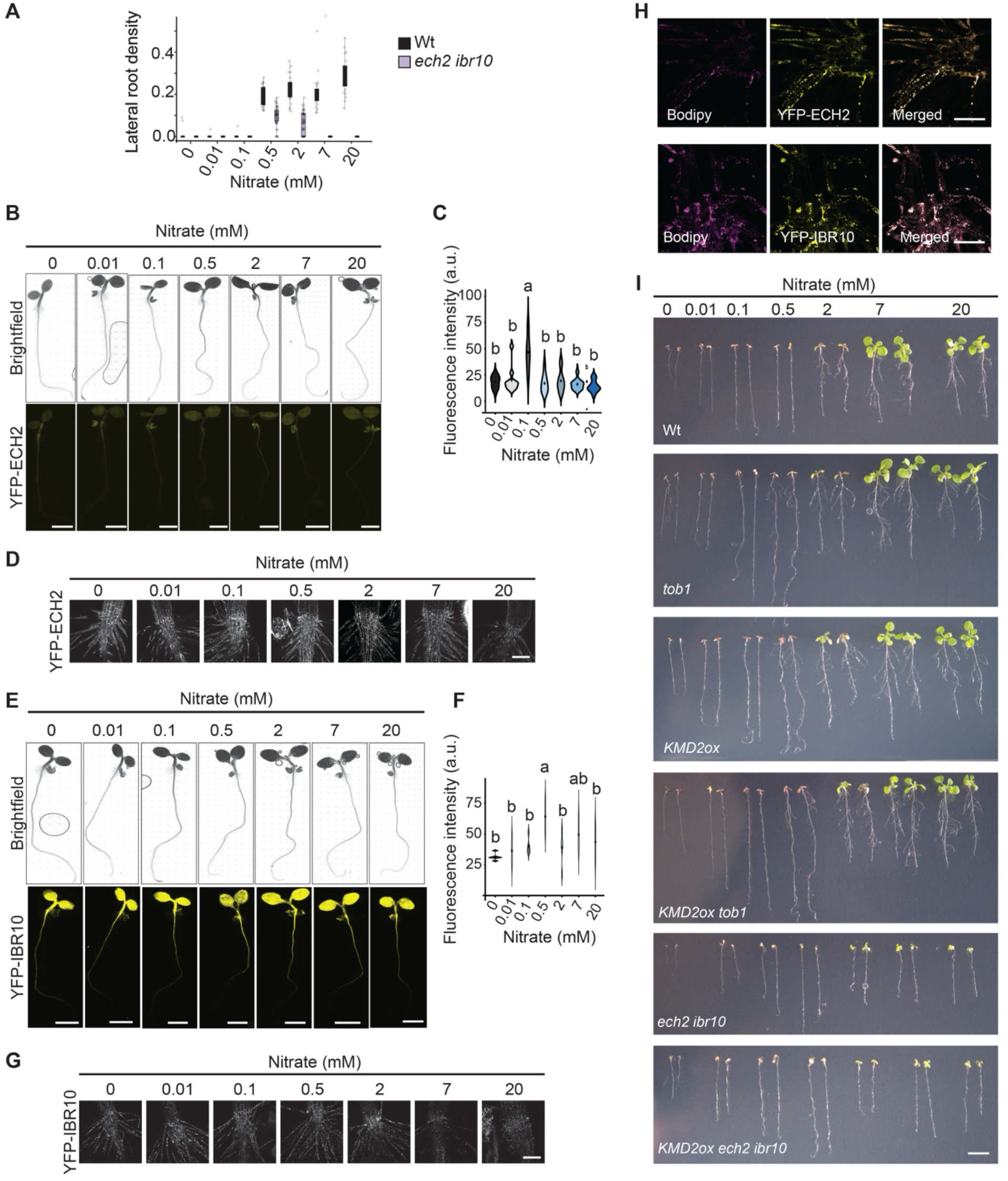
Genes encoding IBA-to IAA-conversion enzymes are expressed in the collet. (A) Nitrate conditions control lateral root density through IBA to IAA conversion. Box plot shows the lateral root density (number of lateral roots per cm of primary root length) of 8-day-old wild type (Col-0) and *ech2 ibr10* seedlings grown in the presence of the indicated nitrate concentrations. Statistical significance between the groups was determined by one-way ANOVA, and significance between each pair was determined using the Wilcoxon signed-rank test (***P ≤ 0.001, **P≤ 0.01, *P ≤ 0.05). Boxes show the first and third quartiles split by mean, and whiskers show the range. Individual data points are shown as jitters. (B) Microscopy images of 4-day-old seedlings expressing *pECH2:YFP-ECH2* germinated on media containing the indicated nitrate concentrations along with the images from Figure 5F for reference. Scale bars = 0.5 cm. (C) Violin plot displaying quantification of fluorescence intensity in collet tissues of *pECH2:YFP-ECH2* expressing 4-day-old seedlings under different nitrate concentrations. n =10 seedlings and statistical significance between the groups was determined by one-way ANOVA; each pair was compared using a Tukey Kramer HSD test and different letters indicate significant differences. (D) YFP-ECH2 expression in the collet under nitrate concentration. Confocal images of pECH2:YFP-ECH2 expression in the collet root hairs of 4-day-old seedling under different nitrate treatments. Scale bars = 250 µM. (E) Microscopy images of 4-day-old seedlings expressing *pIBR10:YFP-IBR10* germinated on media containing the indicated nitrate concentrations along with the images from Figure 5F for reference. Scale bars =0.5 cm. (F) Violin plot displaying quantification of fluorescence intensity in collet tissues of *pIBR10:YFP-IBR10* expressing 4-day-old seedlings under different nitrate concentrations. n =10 seedlings and statistical significance between the groups was determined by one-way ANOVA; each pair was compared using a Tukey Kramer HSD test and different letters indicate significant differences. (G) YFP-IBR10 expression in the collet under nitrate concentration. Confocal images of pIBR10:YFP-IBR10 expression in the collet root hairs of 4-day-old seedling under different nitrate treatments. Scale bars = 250 µM. (H) Colocalization of IBA to IAA conversion enzymes in peroxisomes. Confocal images of collet root hairs stained with bodipy for peroxisome colocalized with YFP-ECH2 and YFP-IBR10. (I) Nitrate-cytokinin-IBA control anchor root formation. Photograph of 8-day-old wild type (Col-0), *tob1, KMD2ox, KMD2ox tob1, ech2 ibr10, KMD2ox ech2 ibr10* seedlings grown under different nitrate concentration. Scale bar = 1 cm.

**Supplemental Figure 5.**
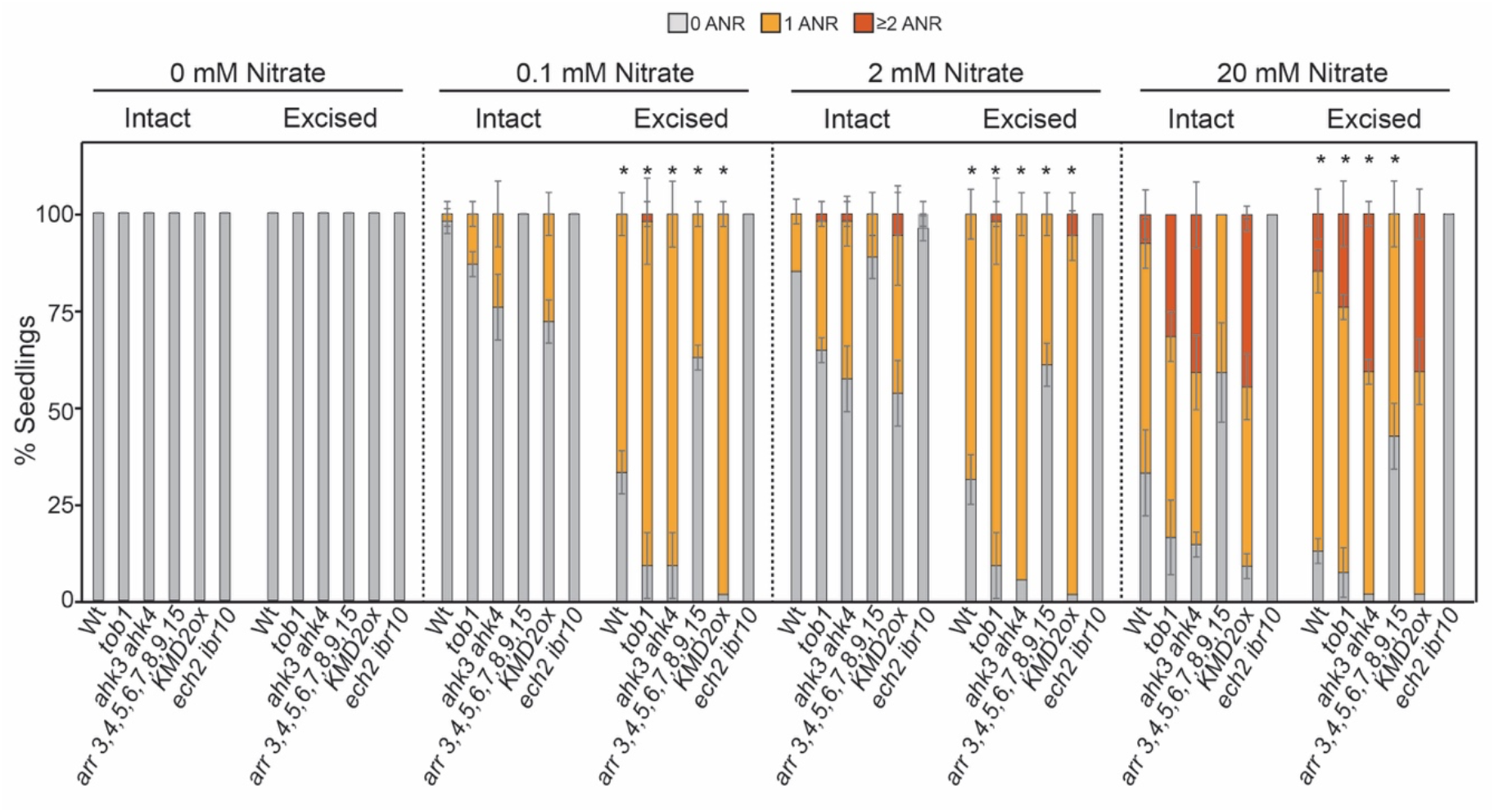
Root tip excision promotes anchor root formation in supraoptimal to high nitrate levels. (A) Excision of root tip promotes more anchor root formation under different nitrate conditions. Quantification of number of anchor roots in wild type, *tob1*, *ahk3 ahk4*, *arr 3,4,5,6,7,8,9,15, KMD2ox* and *ech2 ibr10* seedlings grown in different concentration of nitrate and with intact or excised root tip shown as percentage of seedlings with 0, 1, or ≥2 anchor roots. *n* = 56 and data represents mean ±SD. Statistical significance in anchor root formation between intact and root tip excised seedlings of different genotypes under different nitrate levels were determined by chi-square test.

**TABLE S1.** Composition of plant nutrient (PN) and Hoagland solutions.

| Media | Macronutrient | Micronutrient |
| --- | --- | --- |
| PN (this study) | 2.5 mM $\text{KPO}_4$ | 70 $\mu\text{M}$ $\text{H}_3\text{BO}_3$ |
| | 5 mM $\text{KNO}_3$ | 14 $\mu\text{M}$ $\text{MnCl}_2$ |
| | 2 mM $\text{MgSO}_4$ | 0.5 $\mu\text{M}$ $\text{CuSO}_4$ |
| | 2 M $\text{Ca}(\text{NO}_3)_2$ | 1 $\mu\text{M}$ $\text{ZnSO}_4$ |
| | 40 $\mu\text{M}$ $\text{FeEDTA}$ | 0.2 $\mu\text{M}$ $\text{Na}_2\text{MoO}_4$ |
| | | 10 $\mu\text{M}$ $\text{NaCl}$ |
| | | 0.01 $\mu\text{M}$ $\text{CoCl}$ |
| Hoagland Solution with<br>2.5 mM MES (Jia et al., 2019) | 0.4 mM $\text{K}_2\text{HPO}_4$ | 23 $\mu\text{M}$ $\text{H}_3\text{BO}_3$ |
| | 0.8 mM $\text{MgSO}_4$ | 4.5 $\mu\text{M}$ $\text{MnCl}_2$ |
| | 0.18 mM $\text{FeSO}_4$ | 1.5 $\mu\text{M}$ $\text{ZnCl}_2$ |
| | 5.6 mM $\text{NH}_4\text{NO}_3$ | 0.3 $\mu\text{M}$ $\text{CuSO}_4$ |
| | 0.8 mM $\text{K}_2\text{SO}_4$ | 0.1 $\mu\text{M}$ $\text{Na}_2\text{MoO}_4$ |
| | 0.18 mM $\text{Na}_2\text{EDTA}$ | |

**TABLE S2.**
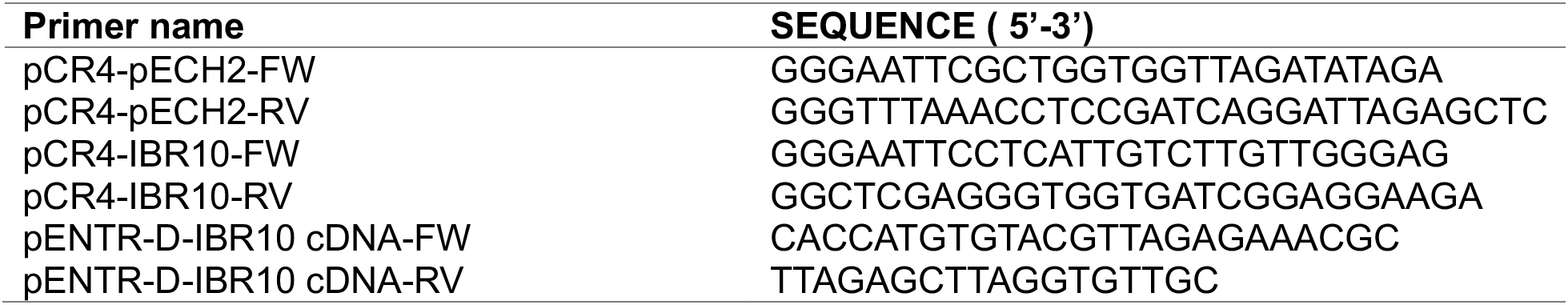
Primers used in this study.

